# No innocent bystanders: pertussis vaccination epitomizes evolutionary parallelisms between *Bordetella parapertussis* and *B. pertussis*

**DOI:** 10.1101/2024.06.18.599646

**Authors:** Valérie Bouchez, Albert Moreno-Mingorance, Alba Mir-Cros, Annie Landier, Nathalie Armatys, Sophie Guillot, Maria Teresa Martín-Gómez, Carla Rodrigues, Julie Toubiana, Ana I. Bento, Michael R. Weigand, Juan José González-López, Sylvain Brisse

## Abstract

Pathogens adapting to the human host and to vaccination-induced immunity may follow parallel evolutionary paths. *Bordetella parapertussis* (*Bpp*) contributes significantly to the burden of whooping cough (pertussis), shares vaccine antigens with *Bordetella pertussis (Bp),* and both pathogens are phylogenetically related and ecological competitors. *Bp* vaccine antigen-coding genes have accumulated variation, including pertactin disruptions, after introduction of acellular vaccines in the 1990s. We aimed to evaluate evolutionary parallelisms in *Bpp*, even though pertussis vaccines were designed against *Bp*.

We investigated the temporal evolution of *Bpp* sublineages, by sequencing 242 *Bpp* isolates collected in France, the USA and Spain between 1937 and 2019, spanning pre-vaccine and two vaccines eras.

We estimated the evolutionary rate of *Bpp* at 2.12×10^−7^ substitutions per site·year^-1^, with a most recent common ancestor of all sequenced isolates around year 1877, and found that pertactin deficiency *in Bpp* was driven by 18 disruptive mutations, including deletion *prn*:ΔG-1895 estimated to have occurred around 1998 and observed in 73.8% (149/202) of post-2007 isolates. In addition, we detected two mutations in the *bvg*A-*fhaB* intergenic region (controlling expression of the master transcriptional regulator BvgA and the filamentous hemagglutinin), that became fixed in the early 1900s.

Our findings suggest early adaptation of *Bpp* to humans through modulation of the *bvgAS* regulon, and a rapid adaptation through the loss of pertactin expression, representing a late evolutionary parallelism concomitant with acellular vaccination against whooping cough.

**IMPORTANCE:** Vaccination against *Bordetella pertussis* (*Bp*) has strongly affected the recent evolution of this main agent of whooping cough. Whether it may have done so co-incidentally on *Bordetella parapertussis* (*Bpp*), which is genetically and ecologically very similar to *Bp,* has not been described in detail. Our findings show striking evolutionary parallelisms of *Bpp* with *Bp*, including early changes in a critical regulatory region, and strong evidence of adaptation to vaccine-driven population immunity, even though whooping cough vaccines were not designed explicitly against *Bpp*. The rapid populational loss of pertactin in countries where acellular pertussis vaccines are used may also reduce protection by vaccination against *Bpp*, the second agent of whooping cough.

## INTRODUCTION

Public health interventions aiming at controlling specific pathogens may concomitantly affect non-target human commensal or pathogenic organisms. For example, commensal bacteria may evolve antimicrobial resistance in response to antimicrobial therapy against pathogens, a phenomenon called bystander evolution [1] that has far-reaching implications in microbial ecology and public health [2]. So far, there is little or no evidence of bystander evolution under vaccination-induced immune pressure. Whooping cough (pertussis) is a human respiratory disease caused mainly by *Bordetella pertussis (Bp).* In 2014, >24 million pertussis cases causing >160,000 deaths in children under 5 years of age were estimated [3]. *Bordetella parapertussis* (*Bpp*) is closely related to *Bp* and also causes whooping cough, though disease is typically less severe [4–6] and thus reported much less frequently than *Bp* infections [7,8]. Further, *Bpp* infection is not reportable in many countries, including the USA. The first whole cell vaccines were already developed using Bp strains when *Bpp* was first reported in 1938 [5], and *Bpp* is still not considered a target of vaccines against whooping cough, which are designed only from *Bp* antigens.

While whole cell pertussis vaccines (wPV), produced using *Bp* strains, remain in use in most of the global South, acellular pertussis vaccines (aPV) were adopted in the mid-1990s and 2000s by many high-income countries. aPVs contain 1 to 5 *Bp* antigens: pertussis toxin (PT), which is always present, combined in most vaccines with filamentous hemagglutinin (FHA), pertactin (PRN) and/or type 2 and type 3 fimbriae (FIM2 and FIM3). It is well established that circulating *Bp* populations, which are human restricted (as is *Bpp*), have evolved in response to vaccine-induced immunity. For example, rates of evolution of vaccine antigen-encoding genes have accelerated since the introduction of aPV, compared to other surface protein genes [9]. Non-synonymous mutations (nsSNP) in PT, PRN, FHA, FIM2 and FIM3 encoding genes, as well as the promoter region of the PT gene cluster, have occurred and raised in frequency, often to fixation in extant *Bp* populations, compared to the pre-vaccine era [10]. Moreover, the rapid emergence of PRN-deficient *Bp* isolates has been observed in countries where aPV vaccination has been implemented [11,12], resulting from multiple independent mutation events rather than the spread of a few genotypes. PRN deficiency has reached near-fixation in early aPV-using countries [10,13], and confers a selective advantage during infection [14,15] that appears higher under the aPV era [16]. The vaccine-driven evolution of *Bp* is regarded as a prominent example of global population-level effects of large-scale vaccination [17,18].

Of the five *Bp* vaccine antigens, *Bpp* expresses orthologs of PRN and FHA only, which are 91.5% and 95.2% identical in their amino acid sequence to their *Bp* counterparts, respectively [19,20]. Given that *Bpp* is phylogenetically related and antigenically similar to *Bp,* the possibility exists that *Bpp* may have also evolved under immune pressure exerted by vaccination targeting *Bp*. PRN-deficient *Bpp* isolates have been observed since 2007 in France [21], but the emergence of PRN-deficient *Bpp* is otherwise undescribed.

*Bp* and *Bpp* have converged in adapting to their human-restricted niche independently, both having evolved from the genetically broader species *Bordetella bronchiseptica* (*Bbs*), an ecological generalist observed in multiple animal host species [22,23]. Among other events, the evolution of *Bp* has involved gene loss, genomic rearrangements and mutations in the intergenic region between the genes coding for FHA and the BvgAS 2-component master regulator of virulence [10,24].

Currently, little is known about the evolution of *Bpp*, largely because it has been rarely isolated in culture. The aim of this study was, by gathering a large international collection of human clinical isolates *of Bpp*, to define its population structure and evolution and explore whether signatures of evolution may be driven by pertussis vaccination-induced immunity.

## METHODS

### Collection of 242 *Bordetella parapertussis* clinical isolates

We collected a large biological resource dataset of *Bpp* isolates from three countries. In France, 119 *Bpp* isolates were collected at the National Reference Center for Whooping Cough and Other Bordetella Infections, isolated between 1937 and 2019. Most of these isolates were collected through the hospital-based pediatric network RENACOQ, which was operated continuously since 1996 [25,26]. From the USA, 85 *Bpp* isolates were collected by Centers for Disease Control and Prevention’s (CDC), being gathered through routine surveillance and during outbreaks between 1937 and 2017, many of which were sequenced as part of a previous study [24]. From Spain, 38 isolates were collected from patients attending the hospitals and primary health care centers according to usual routine diagnostic procedures between 1993 and 2019 in three Spanish regions (Catalonia, Community of Madrid and Castilla-La Mancha). In addition, 8 publicly available genomes of isolates originating from other countries (Australia, Japan, UK and Germany) were included. Details about isolates characteristics are provided in **Table S1**.

### Microbiological characterization, DNA preparation and genomic sequencing

Isolates were grown on Bordet-Gengou agar, antigen characterization was done by Western blot or ELISA, and DNA preparation and genomic sequencing were performed using Illumina technology; details are provided in the supplementary material (Supplementary Method section: Microbiological characterization, DNA preparation and genomic sequencing paragraphs).

### Phylogenetic and gene content analyses

Raw reads were trimmed with Trimmomatic (v. 0.38). Snippy (v. 4.3.6) was used for SNP analysis with *Bpp* strain 12822 (GenBank accession no. BX470249.1) used as reference, without removing recombinant or repetitive regions. A maximum likelihood analysis based on the whole genome SNPs was carried out with IQ-TREE (v. 1.6.10) using 1,000 bootstrap replicates. The existence of a temporal signal in the genomic data was estimated with TempEst by a regression analysis between the root-to-tip divergence in the maximum likelihood tree, and the isolation year (**Fig.S1**). The software tool BEAST version 1.10.4 [27] was used to infer the phylogenetic dynamics as detailed in the supplementary material (Supplementary Method section: Phylogenetic dynamics paragraph).

We looked for transposases using ISMapper [28] or blastN (https://blast.ncbi.nlm.nih.gov/Blast.cgi) with IS*1001* (BPP0078) and IS*1002* (BPP1897) sequences as queries. We estimated the pan genome using Roary version 3.13.0 [29] with default parameters (core genes defined as being present in 95% genomes; without paralogs) from gff3 archives previously annotated with bakta version1.2.1 [30]. We looked for plasmids using PlasmidFinder version 2.1.1 [31].

### Mutations analysis and genotyping

We performed *prn* gene sequence analysis using blastN (https://blast.ncbi.nlm.nih.gov/Blast.cgi) with *Bpp* strain 12822 (NC_002928) gene sequence as queries. Genotyping of virulence genes was done using the BIGSdb platform (https://bigsdb.pasteur.fr/bordetella/) using de novo assemblies as previously detailed [32,33].

## RESULTS

### *Bordetella parapertussis* genomic evolution, population structure and time-resolved phylogeny

We collected 242 *Bpp* isolates between 1937 and 2019 in France, the USA and Spain, conducted genomic sequencing and analyzed the data using phylogenetic and population biology approaches. The number of isolates varied temporally, with a maximum of 32 isolates collected in 2014 (**Fig. 1**). Together with the 8 additional public genome sequences, a dataset of 250 genomic sequences was analyzed.

**Fig. 1.**
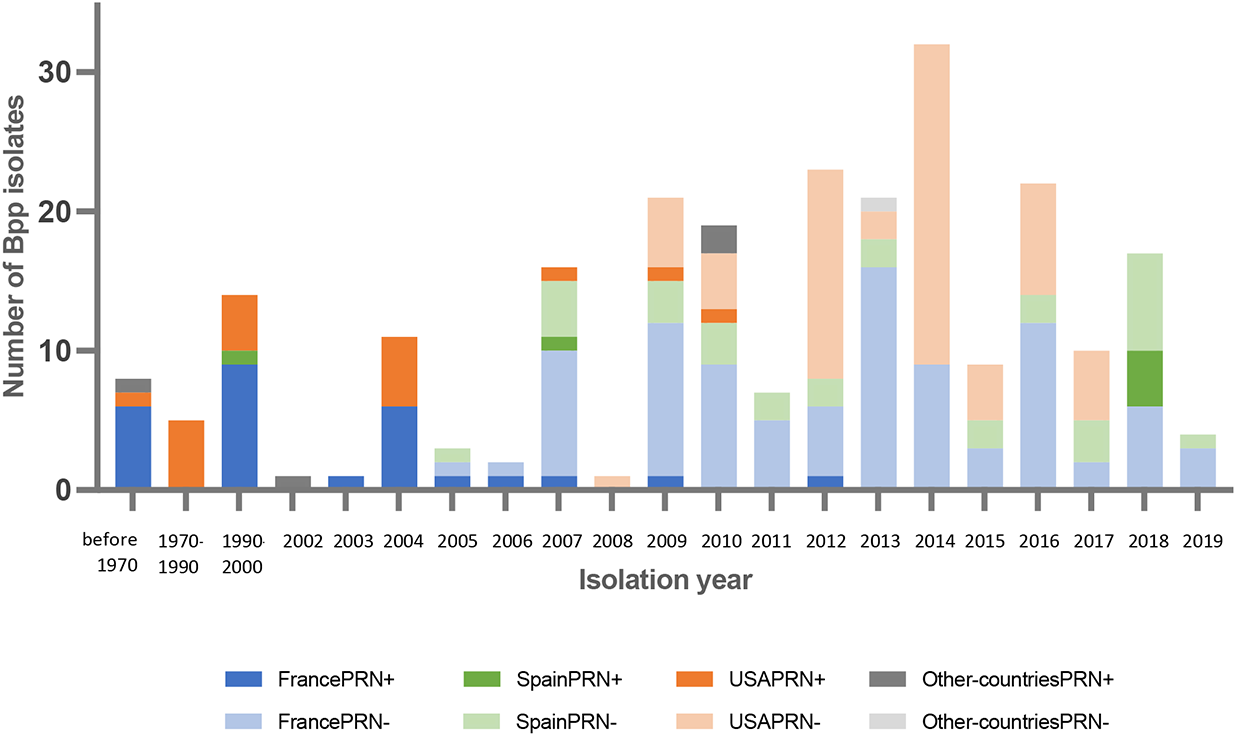
Number of *B. parapertussis* isolates collected per year according to country of origin. This figure includes the 242 collected isolates and 8 isolates for which genomic sequences were publicly available. The data is broken down per year except for the first three bars; Blue: France; Pink: USA; Green: Spain; Grey: others. Dark colors represent PRN-positive (PRN+) isolates and lighter colors PRN-negative (PRN-) isolates (as verified experimentally for France and Spain; and as deduced from genomic sequences for the USA and public sequences from other countries).

The average genome size was 4,732,038 bp and the average G+C% was 68.1%. No plasmid replicons were identified. Using ISMapper, the copy numbers of IS*1001* and IS*1002*, two mobile elements used for diagnosis of *Bpp* [34], were determined to be 22 and 9, respectively, in all isolates except four, which had one or two copies missing. Gene content was highly conserved among the collection of isolates, indicating minimal gene gain or loss. In total, 5,329 different protein-coding genes were identified. Among these, 3,640 were present in at least 99% of isolates, and 4,269 in at least 95%. Genes encoding the main virulence factors and Bvg regulation factors of *Bpp* (*i.e.*, *prn*, *fhaBCD*, *dnt*, *ptlABCDEFGHI*, *ptxAS12345*, *ptxP*, *fim2*, *fim3*, *fimBCD*, *cyaA*, *brkA*, *brkB, bvgA* and *bvgS*) were detected in all isolates except for type IV secretion system encoding gene *ptlD*, where a pseudogene was identified in 9 isolates.

Genome-wide nucleotide variation analysis identified 1,994 single nucleotide polymorphisms (SNP; **Table S2**). On average, strains differed by 47 pairwise SNPs (ranging from 1 to 214). Only 2 mutations were located within IS*1001* transposases. Of the 1,994 SNPs, 35.7% were phylogenetically informative, *i.e.*, the variation was observed in at least 2 genomes. Phylogenetic reconstruction (**Fig. 2A**) uncovered a scaled population structure, within which four main lineages were defined. Lineage 1 comprised 18 isolates placed on early diverging branches of the phylogenetic tree. Lineage 2 corresponded to all isolates (n=23 isolates) with the mutation G3773A within gene *dnt*, leading to the A1258V. A SNP common to lineages 1 and 2 was the A425G nucleotide substitution within gene BPP_RS11415, leading to a N142S change in the corresponding protein. This SNP was absent in isolates from lineages 3 and 4 except for 2 isolates (BBP1_NCBI and B144). Lineage 4 (n=152 isolates) was defined as comprising isolates with the mutation *prn*::delG-1895, which occurred shortly before lineage 4’s MRCA in 1998 [95% HPD: 1996-2001] (**Fig. 2A**). Lineage 3 was defined as comprising the remaining isolates (*i.e.,* with neither the N142S change – with two exceptions – nor the *prn*::delG-1895 mutation). We note that lineages 1 and 2 comprise a previously defined clade 1, whereas lineages 3 and 4 correspond to a previously defined clade 2 [35]. The characteristics of the 4 lineages in terms of pertactin expression are given in the Supplementary appendix. Strikingly, a previously described [24] large genomic rearrangement occurred just before the expansion of lineage 4, in all isolates of which it is observed (**Fig. S7**).

**Fig. 2.**
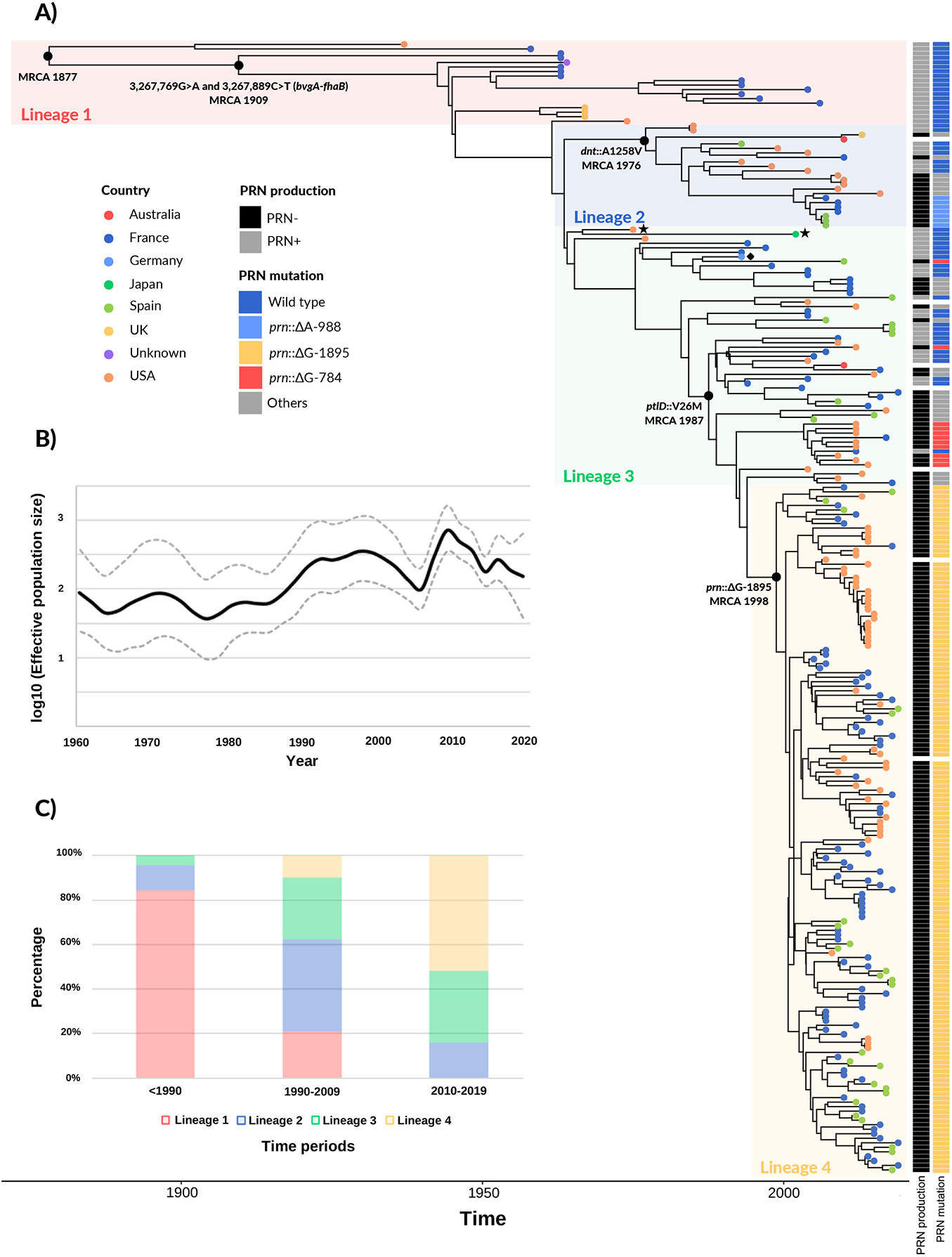
Time-scaled phylogeny of *Bordetella parapertussis*. **Panel A:** Bayesian phylogenetic reconstruction of 250 *B. parapertussis* isolates collected between 1937 and 2019. The phylogenetic tree was built using BEAST (strict clock and Bayesian Skygrid model) from whole-genome SNPs (compared to the reference strain 12822, GenBank accession no. BX470249.1). The reference strain belongs to lineage 3 and its position is indicated by a black rhombus symbol. The two black stars indicate the two lineage 3 isolates with the N142S change (see text). The country of origin of the isolates is represented with colored circles at the tree leaves. Pertactin (PRN) production status, with the three most frequent (>2%) *prn* mutations associated to non-PRN production, are indicated by the two columns on the right of the tree leaves (see color key; see **Table S1** for complete information; missing data are represented in white). **Panel B:** Bayesian Skygrid plot showing temporal changes in effective population size of *B. parapertussis* populations since 1960 (black line) with 95% confidence intervals (discontinuous lines). **Panel C:** proportions of *B. parapertussis* lineages according to three time periods (before 1990, 1990-2009 and 2010-2019). PRN, pertactin; MRCA, most recent common ancestor; UK, United Kingdom; USA, United States of America.

The proportion of the four lineages varied with time: until the mid-1980s, isolates mainly belonged to lineage 1; in contrast after 2010, lineage 4 predominated largely (**Fig. 2B**). We estimated from the genomic data, the fluctuation of the size of the *Bpp* population reflected by our dataset since 1960 **(Fig. 2C)**. The effective population size appeared stable until the mid-1980s, when it began to increase, reaching a maximum in the 2010s. Similar results were obtained when analyzing separately the isolates collected either in the USA, or in Europe (*i.e.,* France and Spain), or when considering random subsamples of 90% of isolates (**Fig. S2**).

A strong temporal signal of SNP accumulation over time was found, with a root-to-tip genetic divergence *versus* time of isolation regression parameter R^2^=0.85 (**Fig. S1**). Bayesian analysis estimated the mean evolutionary rate of *Bpp* as 2.1 x 10^-7^ substitutions per site·year^-1^ (95% highest posterior density [HPD]: 1.9x10^-7^, 2.3 x 10^-7^ substitutions per site·year^-1^), corresponding to 1.02 substitutions per genome·year^-1^. The most recent common ancestor (MRCA) of our *Bpp* dataset was estimated in 1877 [95% HPD: 1865 to 1889]. In turn, the node corresponding to the early diversification of lineages 3 and 2 were estimated to have occurred in 1964 [95% HPD: 1962, 1968] and 1976 [95% HPD: 1974, 1980] respectively, whereas lineage 4 diversification was estimated to have arisen in 1998 [95% HPD: 1996, 2001] (**Fig. 2A**).

### Single nucleotide polymorphisms and insertion and deletion (INDEL) events

Of the 1,994 SNPs, 265 were intergenic and 1,729 were intragenic. Among the latter, 659 were synonymous and 1,070 were non-synonymous (**Table S2)**. SNP densities in genes involved in regulation or coding for hypothetical proteins were statistically higher than the average (p<0.05), whereas SNP densities were statistically lower than average in genes involved in virulence or metabolism (p<0.05) (Supplementary material, SNP densities per functional category paragraph; **Table S3**).

A total of 69 SNPs were located within virulence-associated genes category (**Table S2),** some of which represented landmarks in the evolution of *Bpp*. First, the lineage-2 defining A1258V change in the dermonecrotic factor was inferred to have occurred shortly before 1976 [95% HPD: 1974, 1980]. Second, a SNP observed within gene *ptlD* (leading to a V26M change in the PtlD pertussis toxin export protein) was a marker for a single phylogenetic branch that included part of lineage 3 and the entire lineage 4 isolates; this SNP was observed in 183 isolates (33 of lineage 3 and all of lineage 4) that were collected between 1994 and 2018, and was estimated to have occurred around 1987 [95% HPD: 1984, 1991]. Additional SNPs were also observed in toxins (including the dermonecrotic factor) or in other autotransporters than FHA and pertactin, and in SphB1 (locus tag BPP_RS02120), a serine-protease involved in proteolysis maturation of FHA. Most of these SNPs were observed in only a few *Bpp* isolates. In addition, 18 SNPs were located in genes related to LPS-structural genes and 5 in genes involved in LPS modification (Details are provided in the Supplementary Appendix and in Table S2).

Homoplasic SNPs, *i.e.,* mutations at single positions having occurred in separate branches, are indicative of strong selective pressure leading to convergent evolution. Only 2 of the 1,994 SNPs were homoplasic, including a nonsense mutation leading to a stop codon (Q845Stop) in *prn*, observed in two isolates from lineages 2 and 3 (see Supplementary Results section).

Regarding small insertions and deletions, 329 INDEL events (as compared to the genome sequence of reference isolate 12822) were observed. Only 29 INDEL events were present in more than 10 isolates; 20 of these were located out of coding regions, and 9 within them (see INDEL, **Table S4**). Of these, 7 were inferred to induce frameshifts, including 2 INDELs within the *prn* gene and one within *bscR*. Two INDELS were also observed within *fhaB,* but only affected 3 and 1 isolate, respectively (**Table S4**). As described above, the single G deletion in *prn* position 1895 (*prn*::ΔG-1895) was present in 150/152 from lineage4 (for the H299 and H602, the *prn* sequence did not allow to confirm the presence of the mutation).

### Pertactin gene diversity and its multiple disruptions

Only 51 of 250 isolates had a *prn* nucleotide sequence (locus tag BPP_RS05740) identical to the reference strain 12822, including all isolates of lineage 1 (n=18 isolates) and some isolates of lineage 2 (n=8) and lineage 3 (n=25, including the reference). PRN production was confirmed experimentally for 27 of these 51 isolates (**Table S1**).

For 192 of the 199 remaining isolates, the *prn* sequence had a point mutation, frameshift, or insertion sequence mutation (**Table S1)**. We found a total of 18 distinct mutations. Four mutations were nonsense SNPs resulting in stop codons, whereas 13 were insertions or deletions events inferred to lead to *prn* deficiency (**Table S5**). These distinct *prn* mutations were scattered in lineages 2, 3 and 4 and one of them (described above) was homoplasic (**Fig. 2**). Last, *prn*::ΔG-1895 deletion was by far the most frequently observed mutation, having been associated with the expansion of lineage 4. PRN production was confirmed experimentally to be deficient for 125 of the 192 isolates corresponding to all the different types of mutations (**Table S1**). Overall, 56.5% (13/23) of lineage 2 isolates, 50.9% (29/57) of lineage 3 isolates and 98.7% (150/152) of lineage 4 isolates were demonstrated or inferred to be PRN deficient based on the presence of one of the 18 *prn* mutations.

Whereas region 1 of PRN is highly variable in *Bp* [36], almost all *Bpp* isolates with a full length gene displayed the same number of repeats in the two repeat regions of *prn* (4 repeats in region 1 and 9 in region 2), consistent with previous reports [37,38].

### SNPs in the *fhaB-bvgA* intergenic region, and in FHA and functionally related genes

An exceptionally high SNP density was observed in the intergenic region located between *fhaB* and *bvgA,* with six intergenic SNPs **(Fig. 3**) (**Table S3**). Whereas four SNPs located in the phosphorylated BvgA binding site just upstream of *fhaB* gene (also corresponding to P3 bvgA promoter) were each observed in a few isolates (all collected after 2004), two SNPs were largely shared by *Bpp* isolates, located at position 3,267,769 (G>A) and 3,267,889 (C>T). Both occurred in an early branch of the phylogeny (**Fig. 2**), with an MRCA estimated in 1909 (95% HPD: 1899, 1921). These two mutations have been fixed in extant *Bpp* populations, as isolates that did not carry these SNPs were not observed after 1958. While the first of these SNPs is located 23 nucleotides upstream of the -35 box of *fhaB*, the SNP at position 3,267,889 (C>T) is located within the -35 element of *bvg*A gene, changing the element from TT**C**AGAA to TT**G**AGAA, clearly suggesting an impact on gene expression (**Fig. 3**).

**Fig. 3.**
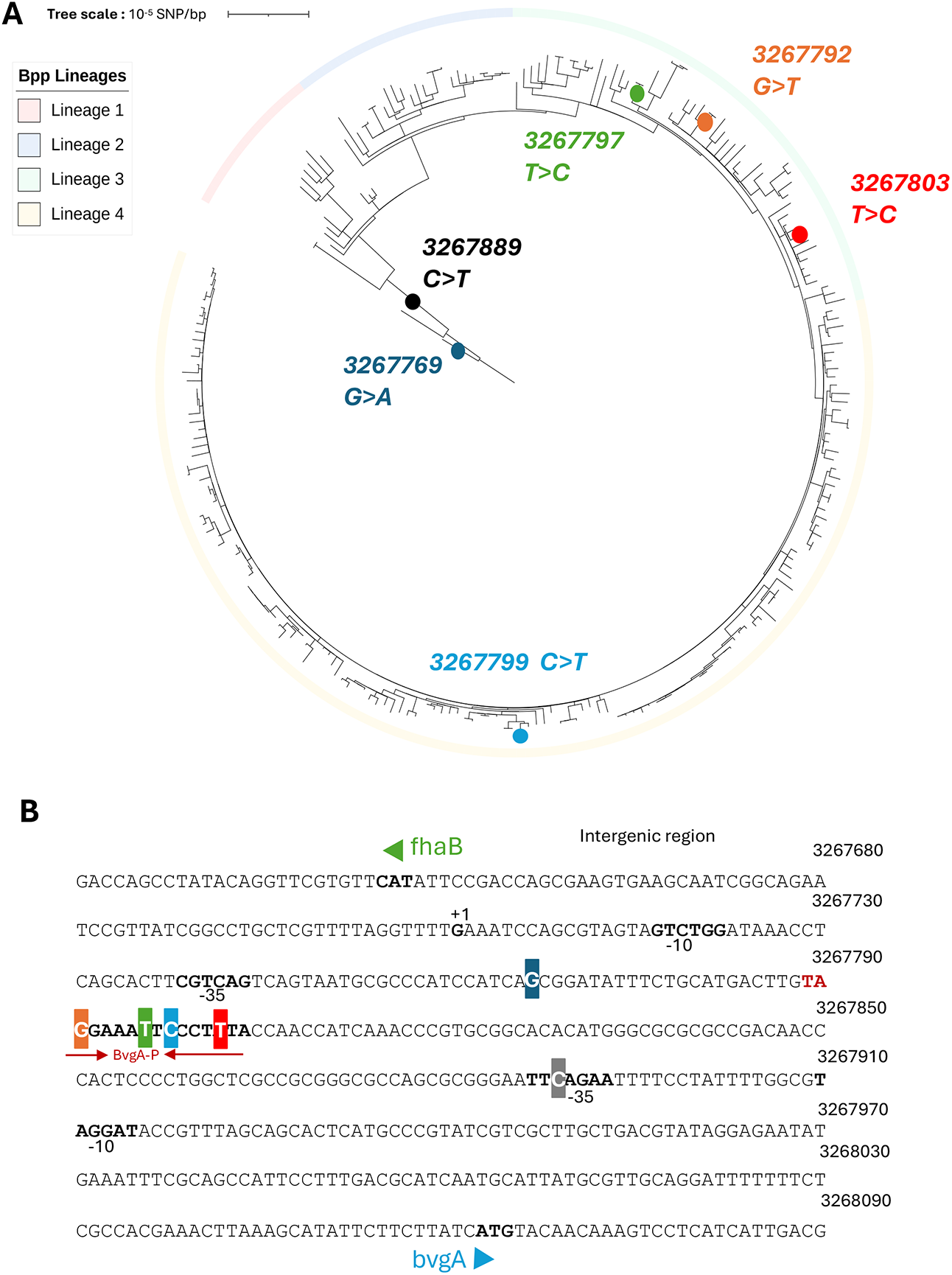
Mutations located in the intergenic region between *fhaB* and *bvgA*. **Panel A:** Circular phylogenic tree based on SNPs, rooted on isolate Bpp63.34. The four lineages are represented with colored branches as in Figure 2. Lineage 1: light red; Lineage 2: light blue; Lineage 3: light green; Lineage 4: light orange. Mutations observed within the *fhaB-bvgA* intergenic region are indicated by circles on tree branches and labelled with their nucleotide position in the genome. **Panel B:** Precise localization of the observed mutations (highlighted with color background as in panel A) within the intergenic sequence. The minus 10 and minus 35 motives upstream of both genes, the +1 transcription start site of *fhaB*, and the BvgA binding motif (from 3267789 to 3267804), are indicated in bold.

FHA production was confirmed experimentally for 145/250 isolates of the study, as evidenced using either Western blots or ELISA (**Table S1**). We nevertheless observed 11 SNPs within the *fhaB* gene itself (locus tag BPP_RS15295), all being found in only one or a few isolates (**Table S2**). Of these, six were non-synonymous, two of which affecting the mature FHA [31]: one at position 3,261,750 (V1963A), observed in four isolates of lineage 1; and one at position 3,266,524 (T372A), observed in a single isolate (J324) of lineage 3 (**Table S2**).

nsSNPs were also observed affecting proteins functionally related to FHA, including FhaS, FhaJ and SphB1, a serine-protease involved in proteolysis maturation of FHA [39] (**Table S2;** detailed in supplementary appendix).

## DISCUSSION

Acellular vaccines against *B. pertussis* (*Bp*) are in use since more than 20 years in multiple countries including the USA, France and Spain. Here, we addressed the question of the possible impact that large-scale whooping cough vaccination might have exerted on the second agent of whooping cough, *B. parapertussis* (*Bpp*), even though this organism was not the explicit target of the vaccine. Because *Bpp* expresses two antigens, FHA and pertactin, that are closely related orthologs of *Bp* vaccine antigens and are part of the *bvgAS* virulence regulon, changes in these proteins may have important consequences on the current epidemiology and pathogenesis of *Bpp* infections.

By taking advantage of a unique dataset of human isolates collected over 83 years in three different countries, we also address the broader biological questions of the genomic diversity, population structure and genome-scale evolution of the so-far elusive *Bpp,* and the possible evolutionary parallelisms that this pathogen might show with *Bp*, its close relative and ecological competitor. Our data reveal, over time, the successive replacement of *Bpp* subpopulations by more recently emerged ones. By considering four deep *Bpp* phylogenetic lineages, we showed how their relative proportions have shifted: whereas lineage 1 was predominant before 1990, lineage 4 became the most frequent since 2010, now being nearly exclusive. This temporal replacement pattern is reminiscent of the disappearance of ancient lineages of *Bp,* which were replaced among extant infectious isolates by more recently evolved sublineages [10]. This scaled phylogenetic structure pattern is also typically encountered in human viruses evolving to escape previously built host immunity by antigenic drift, such as Influenza virus [40] or more recently SARS-CoV-2 [41]. The expansion of lineage 4, characterized by a genomic rearrangement and the lack of PRN production, coincides with the peak detected in the effective population size analysis, around the year 2010, suggesting that isolates from lineage 4 may have a better fitness in the three surveyed countries using acellular vaccines. The evolution of *Bpp* by successive lineage replacement might be driven by a combination of its ongoing adaptation to humans, natural immunity built in human populations as a result of infection, ecological interactions with *Bp* [42,43] and possibly also by vaccination-induced immunity, which we discuss below.

The *Bpp* population genomics data also revealed a regular pattern of SNP accumulation over time, enabling us to estimate the mutation rate of *Bpp* at 2.1x10^-7^ substitutions per site·year^-1^. This rate is remarkably similar to the one estimated within the main branch of *Bp* (2.24x10^-7^ substitutions per site·year^-1^) [10], consistent with the shared recent ancestry and similar ecology of these two pathogens, which both evolved from their progenitor genomic species *B. bronchiseptica* [10,22,33]. Remarkably, the estimated most recent common ancestor of human *Bpp*, around year 1877, is more recent by only a few decades than the main *Bp* branch, which was estimated to have emerged between years 1790 and 1810 [10]. The crowding and promiscuity that increased rapidly during the industrial revolution in the 19^th^ century represents a possible driver of the expansions of *Bpp* and *Bp* in the populations of developed countries. Clearly, our sample (mainly from three Northern hemisphere countries) may miss deeper branching isolates that could be circulating elsewhere or have become extremely rare, similar to the exceptionally rare deep lineage of *Bp* [10]. Note that ovine *Bpp* isolates, which were seldom reported and for which only a single genomic sequence is available [37,44,45], were not considered in this study because they belong to a separate evolutionary lineage [22] and hence would not affect the above temporal analyses and conclusions.

FHA is one of the components of most (but not all) current aPVs, and BvgA is the main regulator of virulence genes in *Bbs* and *Bp.* The *bvgA-fhaB* intergenic region was shown to have undergone extensive evolution in *Bp* [10], which may impact not only the expression of both genes, but also those of the *bvgAS* regulon [46]. Here we report several mutations located in the orthologous intergenic region of *Bpp*. One of these mutations (in position 3,267,792 in **Fig. 3** or 155 in **Fig. S3**) is at the exact same position as a mutation observed in *Bp* [10,34]. Although six SNPs were observed in the *bvgA-fhaB* intergenic region of *Bpp*, two of these occurred early in the evolutionary history of this pathogen (estimated around 1909) and became fixed in *Bpp*. These two mutations predate largely pertussis vaccination and may have been selected for adaptation to humans, which were recent novel hosts for *Bpp* at that time, or in reaction to natural infection-driven immunity. The high SNP density in the *bvgA-fhaB* intergenic region in both *Bp* and *Bpp* suggests a central role of this critical regulatory region in evolutionary adaptation to the human niche, as both pathogens diverged from their ecological generalist progenitor species *B. bronchiseptica*. Whether and how these two ancestral *Bpp* intergenic mutations have impacted the levels of *fhaB* expression, *bvgA* expression, or both, thus stands out as a central question to decipher the adaptive trajectory of *Bpp*.

The four other *bvgA-fhaB* intergenic SNPs are located in the phosphorylated BvgA binding site of the *fhaB* promoter [10,46], also suggests a functional impact of these mutations, but as these were observed in only a few isolates, they may reflect a transient selective advantage, perhaps in patients with atypical anti-FHA immunity. Further non-synonymous genetic variation in *fhaB* and functionally related genes was observed (supplementary Appendix). Mutations in the coding sequence of *fhaB* may reflect the fine-tuning of FHA protein interaction with its receptor, even though none were located in the FHA-RGD motif. Several mutations also occurred in *Bp* within the gene encoding FHA [10,16]. We confirmed FHA production experimentally in all (145/250) tested *Bpp*, and as FHA-negative *Bp* are exceptional too [25], this protein seems to exert an essential role in the biology of both agents of whooping cough. Overall, these observations point to a particular role of FHA and related functions in *Bpp* biology, as in *Bp* [47].

Regarding the effect of whooping cough vaccination on *Bpp*, our work uncovers a number of genetic signatures of evolution in the genes coding for the two *Bp* vaccine antigens which are expressed by *Bpp*. As *Bpp* produces neither pertussis toxin, due to several mutations within the promoter sequences of the synthesis gene cluster [19,48], nor the fimbriae FIM2 and FIM3 proteins (even though their genes are present and undisrupted in *Bpp* genomes) [20,34], the lack of evidence for selection in these other antigens acts as an interesting control. No SNP was observed within fimbriae genes: neither within *fimABCD* structural genes nor within *fimX* or *fimN* genes, which code for other fimbriae subunits. This lack of variation is consistent with the lack of expression of these genes in *Bpp* [20], which implies an absence of positive selection to optimize interactions with host receptors and to escape immunity. Similarly, as pertussis toxin is also not produced by *Bpp*, the mutations we observed may be considered as contributing further to the gene decay of the pertussis toxin gene cluster.

In *Bp*, pertactin deficiency is a major recent evolutionary phenomenon, shown to be driven mainly by acellular vaccine (aPV)-induced immunity [12,39,49]. But so far, the genetic evolution of pertactin expression in *Bpp* has been little documented [36,50]. Our data provide strong evidence for the evolution of this antigen being driven by acellular pertussis vaccines too. First, we observed a population shift towards pertactin deficiency in *Bpp*, which has started just after the roll-out of aPVs. Second, besides the prominent *prn*::ΔG-1895 mutation, 17 other pertactin deficiency mutations were identified, and all were dated between 2005 and 2018. This convergent pattern of gene disruption after the introduction of aPV strongly supports the view that pertactin expression by *Bpp* is disadvantageous in aPV countries. Data from three countries that use wPV instead of aPV further support this hypothesis, as no pertactin-deficient was observed in *Bpp* isolates collected between 1998 and 2015 [35,51,52]. Although more *Bpp* sampling would strengthen the trend we observed, the pertactin-deficient population increase seems to be even faster in *Bpp* than observed for *Bp* **(Fig. S5),** as almost all *Bpp,* but only 50% to 90% *Bp*, depending on country, are now deficient [53,54].

Thus, even though pertussis vaccines were designed against *Bp*, the main whooping cough agent, our genomic analyses indicate that *Bpp* has indeed been affected by pertussis vaccination. Although the phylogenetic proximity and shared antigens of *Bp* and *Bpp* makes the ‘bystander’ status of *Bpp* questionable, our work uncovers a clear evolutionary impact of vaccination on an organism that was not the explicit target of the vaccine. In the strict sense, this work thus demonstrates a bystander impact of vaccination on a non-target organism.

Vaccination against *Bp* has been considered to have low, or even no, efficacy against *Bpp* [55–57]. The strong evidence provided here of aVP vaccination driving pertactin deficiency in the populations of *Bpp,* indeed suggests a cross-protection of aPV pertussis vaccines on *Bpp* isolates that produce pertactin [12], which we hypothesize to exert the selective disadvantage we observed in the three aVP vaccinated populations that we surveyed. An important implication is that, as extant isolates of *Bpp* now rarely produce pertactin, cross-protection against *Bpp* from whooping cough vaccines may have weakened significantly in the last 20 years. This evolution leaves only FHA as an aVP vaccine antigen expressed by *Bpp*, against which the bactericidal activity of antibodies is weak [49]. Future improved whooping cough vaccines could benefit from comprising *Bp-Bpp* cross-reacting antigens explicitly, such as the adenylate cyclase [58] or conserved antigens identified through immune-informatics [59], or could incorporate *Bpp*-specific antigens, such as the O-antigen [60].

In conclusion, our study provides important novel insights into the past evolutionary dynamics of *Bpp* and uncovers a remarkable picture of parallel evolution between the adaptation of *Bpp* and *Bp* populations to humans, including their timing of emergence, rate of evolution, successive lineage replacement, early adaptation to the human niche, and vaccine-driven evolution. These parallelisms illuminate how two distinct pathogens that have evolved from a single common ancestral species, have adapted to the human host and later in response to vaccination-induced immunity. The deep evolutionary picture we uncovered for *Bpp*, highlights the bystander effect of pertussis vaccination against *Bpp* as the latest example of the evolutionary parallelism between the two agents of whooping cough.

## Acknowledgements

We are grateful to Nicole Guiso, who directed the French NRC between 1994 and 2016 and contributed to the initiation of the French whooping cough surveillance network (RENACOQ). We also acknowledge the contributions to this sentinel surveillance network, of Emmanuel Belchior, Fatima Aït Belghiti and Daniel Levy-Bruhl from Public Health France (Santé Publique France), and thank the French microbiologists who sent clinical isolates: Nathalie Brieu (Aix-en-Provence), Farida Hamdad (Amiens), Marie Kempf and Hélène Pailhoriès (Angers), Cécile Jensen (Avignon), Philippe Lehours and Jennifer Guiraud (Bordeaux), Hervé Le Bars (Brest), Christophe Isnard (Caen), Nathalie Wilhelm and Alain Le Coustumier (Cahors), Julien Delmas (Clermont-Ferrand), Dominique De Briel and Laurent Souply (Colmar), Saïd Aberrane (Créteil), Marie Coudé (Le Mans), Fabien Garnier (Limoges), Ghislaine Descours (Lyon), Hélène Jean-Pierre (Montpellier), Corentine Alauzet (Nancy), Sophie-Anne Gibaud (Nantes), Stéphane Bonacorsi (Paris, Hôpital Robert Debré), Lucien Brasme (Reims), Ludovic Lemée (Rouen), Christelle Koebel (Strasbourg), Philippe Lanotte (Tours), Stéphane Bland and Hélène Petitprez (Annecy), Didier Raffenot and Marion Levast (Chambéry), Florence Doucet-Populaire and Nadège Bourgeois-Nicolaos (Clamart), Christophe Burucoa (Poitiers), Florence Grattard (Saint-Etienne), Stéphanie Marque-Juillet (Versailles).

We further thank the technical staff and microbiologists of the Clinical Microbiology Laboratory of Hospital Universitari Vall d’Hebron and the participants of FIS PI18/00703 project: Raquel Abad (Centro Nacional de Microbiología), M. Ángeles Orellana (Hospital Universitario 12 de Octubre), Iván Bloise and Manuela de Pablos (Hospital Universitario La Paz), M. Elena Rodríguez and Alejandro González-Praetorius (Hospital Universitario de Guadalajara), M. Nieves Gutiérrez (Complejo Asistencial Universitario de Salamanca), Carmen Muñoz (Hospital Sant Joan de Déu Barcelona), Magda Campins and Sonia Uriona (Hospital Universitari Vall d’Hebron), Carlos Rodrigo (Hospital Universitari Germans Trias i Pujol), M. José Vidal (Agència de Salut Pública de Catalunya).

We thank Lucia Tondella, Yanhui Peng, and Matthew Cole (Centers for Disease Control and Prevention) as well as the Enhanced Pertussis Surveillance/Emerging Infections Program Network sites and other US state public health departments for contributing isolates. The findings and conclusions in this report are those of the authors and do not necessarily represent the official position of the Centers for Disease Control and Prevention.

## Authors license statement

This research was funded, in whole or in part, by Institut Pasteur and Santé publique France. For the purpose of open access, the authors have applied a CC-BY public copyright license to any Author Manuscript version arising from this submission.

## Ethical statements

All French bacteriological samples and associated clinical data are collected, coded, shipped, managed and analyzed according to the National Reference Center protocols that received approval by French supervisory ethics authority (CNIL, n°1474593). The study of the Spanish isolates was approved by the Ethics Committee of the Vall d’Hebron Hospital (reference number: PR(AG)694/2020). This activity was reviewed by CDC, deemed not research, and was conducted consistent with applicable federal law and CDC policy. See e.g., 45 C.F.R. part 46, 21 C.F.R. part 56; 42 U.S.C. §241(d); 5 U.S.C. §552a; 44 U.S.C. §3501 et seq.

## Author’s contributions

Valérie Bouchez gathered WGS data, analyzed SNP data and PRN genotypes, and wrote the initial versions of the manuscript together with Sylvain Brisse. Annie Landier and Nathalie Armatys performed the experimental work on French isolates, supervised by Sophie Guillot and Carla Rodrigues, who validated the data. Maria Teresa Martín-Gómez performed the laboratory work and participated in the characterization of the Spanish isolates. Alba Mir-Cros and Albert Moreno-Mingorance gathered WGS, performed SNP and Bayesian analyses and wrote initial parts of the manuscript. Ana Bento supervised the Bayesian analyses. Michael Weigand provided WGS from USA and analyzed the genomic rearrangements data. Julie Toubiana provided input for the collection and validation of the clinical data. Juan José González-López and Sylvain Brisse conceived and coordinated the study. All authors revised and agreed on the last version of the manuscript.

## Data availability

The genome sequence reads were deposited in European Nucleotide Archive and are available from accession number PRJEB45017, and in the National Center for Biotechnology Information accession number PRJNA731630 (**Table S1**).

